# The Polarization Dependence of 2D IR Cross-Peaks Distinguishes Parallel-Stranded and Antiparallel-Stranded DNA G-quadruplexes

**DOI:** 10.1101/2021.07.13.452271

**Authors:** David A. Price, Poornima Wedamulla, Tayler D. Hill, Taylor M. Loth, Sean D. Moran

## Abstract

Guanine-rich nucleic acid sequences have a tendency to form four-stranded non-canonical motifs known as G-quadruplexes. These motifs may adopt a wide range of structures characterized by size, strand orientation, guanine base conformation, and fold topology. Using three K^+^-bound model systems, we show that vibrational coupling between guanine C6=O and ring modes varies between parallel-stranded and antiparallel-stranded G-quadruplexes, and that such structures can be distinguished by comparison of polarization dependent cross-peaks in their two-dimensional infrared (2D IR) spectra. Combined with previously defined vibrational frequency trends, this analysis reveals key features of a 30-nucleotide unimolecular variant of the Bcl-2 proximal promoter that are consistent with its reported structure. This study shows that 2D IR spectroscopy is a convenient method for analyzing G-quadruplex structures that can be applied to complex sequences where traditional high-resolution methods are limited by solubility and disorder.

## 1. Introduction

G-quadruplexes are non-canonical four stranded secondary structure motifs formed by guanine-rich nucleic acid sequences.*^1^* They have been implicated in a wide variety of biological processes, such as DNA replication, the regulation of gene expression, and telomere maintenance, and may also play a role in neurodegenerative diseases.*^1,2^* The core structure of a G-quadruplex is a stack of planar G-quartets, which are stabilized by a combination of Watson Crick and Hoogsteen hydrogen-bonding as well as metal ion coordination to guanine C6=O carbonyls.*^3^* Despite this commonality, G-quadruplexes form a diverse family of structures that vary in the number of G-quartet layers, conformations of individual guanine residues, strand orientations, and topologies.*^3,4^* The ability to distinguish the features of a G-quadruplex fold is central to understanding their structure-function relationships, as well as in efforts to design molecular probes and therapeutics that target them.*^4–6^* A number of biophysical techniques have been employed to do so; among them, high-resolution NMR and X-ray methods provide the most detailed structural information.*^7^* Recently, NMR ^13^C and ^1^H chemical shifts have also been employed to identify guanine base conformations (*syn*- or *anti*-) and fold topologies from the torsional angles of glycosidic bonds.*^8^* However, both NMR and crystallographic approaches are limited by sample uniformity and solubility,*^4^* and many biologically relevant G-quadruplex forming sequences are disordered or have a tendency to aggregate. In such cases, other methods including UV circular dichroism (UV-CD),*^9^* fluorescence emission,*^6,10^* mass spectrometry,*^11^* and vibrational spectroscopies*^12^* are valuable for assigning G-quadruplexes to topological classes.

Infrared methods are sensitive to molecular structures because vibrational frequencies and lineshapes depend on local electrostatics and coupling between oscillators. Ultrafast two-dimensional infrared (2D IR) spectroscopy is a powerful probe of these phenomena due to its ability to observe dynamics via spectral diffusion, to resolve peaks in congested spectra, to measure the anharmonicites of vibrational modes, and to directly detect coupling through the appearance of cross-peaks.*^13,14^* In peptides and proteins, these advantages provide detailed insight into secondary structure compositions via the analysis of amide I modes.*^15–17^* In particular, extended arrays of coupled chromophores in amyloid ß-sheets produce distinctive spectral features that can be used to assess differences in molecular structures between systems.*^18–22^* In nucleic acids, 2D IR spectroscopy has been applied to study the hydration and ionic interactions of phosphate groups, the structures and dynamics of base pairs, and base stacking.*^14,23–28^* Precedent from the amyloid literature suggests that 2D IR spectra may be used to distinguish between topological variants of G-quadruplexes, which also comprise strongly coupled arrays of chemically identical chromophores.

Upon the formation of a G-quadruplex, characteristic frequency and lineshape changes are observed among four anharmonically-coupled modes of the guanine base (denoted here as guanine I-IV) that appear in the region of 1500 – 1700 cm^−1^.*^29^* Previous Fourier transform infrared (FTIR) studies suggested that the magnitudes of frequency shifts may distinguish parallel-stranded and antiparallel-stranded structures.*^12^* Recently, we used 2D IR spectroscopy and heavy isotope labeling to demonstrate that guanine C6=O (guanine I) frequency shifts in parallel-stranded G-quadruplexes, which adopt all-*anti*-base conformations, arise from vibrational coupling between stacked G-quartets and scale with the number of consecutive ordered layers.*^27^* This finding is consistent with smaller guanine I blue shifts in antiparallel-stranded G-quadruplexes because reorientation of transition moments in their mixed *syn*-/*anti*-structures attenuates coupling constants.*^30^* However, it also indicates that the effects of motif size and conformation are impossible to quantify from frequencies alone. In this Article, we demonstrate that variations in guanine base conformations between parallel-stranded and antiparallel-stranded G-quadruplexes result in a polarization dependence of 2D IR cross-peak intensities that is a convenient spectral marker of strand orientations and topologies. By combining this information with known frequency trends, we determine key structural features of a G-quadruplex formed by a 30-nucleotide variant of the Bcl-2 promoter.

## 2. Materials and Methods

### 2.1. Molecular modeling

Models of K^+^-bound G-quadruplexes were prepared by substituting K^+^ into Na^+^-bound or vacant sites in their respective PDB structures (PDB IDs: 244D, 2GWQ, and 201D, respectively). After substitution, the ion positions were relaxed via minimization using the AMBER10:EHT force field in MOE (CCG Inc., Montreal, Quebec) with all other atoms fixed. A second unconstrained minimization step using the same force field was used to relax the complexes.

### 2.2. Sample preparation

D_2_O (99.9%) was purchased from Cambridge Isotope Laboratories (Tewksbury, MA). All other chemicals were obtained from Sigma-Aldrich (St. Louis, MO) and used as received. Oligonucleotides were obtained from Integrated DNA Technologies (Coralville, IA). G-quadruplexes were assembled according to methods described previously.*^27^* Briefly, oligonucleotides were dissolved in 20 mM tris buffered D_2_O (pD 7.5), supplemented with the desired concentration of KCl, to a final concentration of either 3 mM (model systems) or 1 mM (Bcl-2 promoter). After dissolution, the samples incubated overnight at 4 °C to deuterate exchangeable sites. Solutions were then heated to 90 °C for 10 minutes and allowed to cool slowly to room temperature for analysis. Guanosine monophosphate (GMP) samples were prepared at 50 mM concentration in the same manner, but KCl was omitted from the buffer solutions.

### 2.3. Native PAGE

Model G-quadruplexes prepared with 140 mM KCl were loaded onto an 18% 19:1 acrylamide:bis-acrylamide gel and run at 20 °C using 20 mM tris (pH 7.2) with 40 mM KCl as a running buffer. A marker composed of a mixture of single-stranded polyT oligonucleotides was included as a reference. The gel was fixed at 4 °C using 50% ethanol, 37% H_2_O, 10% glacial acetic acid, 3% glutaraldehyde, and was stained with methylene blue for visualization.

### 2.4. 2D IR spectroscopy

2D IR spectra were collected using a partially collinear 2DQuick Array pulse-shaping spectrometer (PhaseTech Spectroscopy, Madison, WI) equipped with a 128 x 128 pixel MCT array detector. Mid-IR pulses (~17 μJ) centered at 6 μm were produced using by pumping a TOPAS-Prime optical parametric amplifier equipped with a AgGaS_2_ difference frequency mixer (Light Conversion, Vilnius, Lithuania) using 1.75 mJ pulses (100 fs, 790 nm) from an UpTek Solutions (Bohemia, NY) Phidia Ti:Sapphire amplifier.

For data collection, 1 μL of each sample was placed between two 2 mm CaF_2_ windows separated by a 50 μm PTFE spacer in a demountable transmission cell. All spectra were collected at 20 °C. The sample compartment was continuously purged with dry air to eliminate interference from water vapor. The relative polarization of the pump pulse pair (k_1_, k_2_) to the probe (k_3_, LO) was set by rotating a λ/2 waveplate and linear polarizer after the pulse shaper. In our notation *X* and *Y* correspond to orthogonal linear polarizations of the pulses, and *Z* is normal to the sample plane. All spectra were collected with a pump-probe waiting time set to 0.0 ps relative to the maximum intensity of dimethylformamide dissolved in D_2_O. Probe frequencies were calibrated using the absorbances of 4-nitrobenzaldehyde and 2-hydroxy-5-nitrobenzaldehyde dissolved in dichloromethane as external standards. Time-domain data was averaged, apodized using the Hamming window function, and Fourier transformed in MATLAB (MathWorks, Natick, MA) to generate the 2D IR spectra.

### 2.5. FTIR spectroscopy

FTIR spectra, were collected using a JASCO 6800 FTIR spectrometer equipped with a single-channel MCT detector and a PIKE (Madison, WI) MIRacle ATR accessory with a ZnSe crystal. Samples were placed between two CaF_2_ windows in a demountable cell as described above. The optical compartment was purged continuously with dry air to eliminate interference from water vapor. Buffer backgrounds were subtracted from spectra and residual baseline correction was performed using low-order polynomial fitting in MATLAB.

## 3. Results and Discussion

### 3.1. Characterization of model G-quadruplexes

To examine the influence of guanine base conformations on the signatures of vibrational coupling in G-quadruplexes, we selected three DNA oligonucleotides based on the *Oxytrichia nova* telomeric repeat (Table S1) that form stable K^+^-bound complexes composed of four stacked G-quartets. Models of the potassium-bound structures are shown in Fig. 1A for (**1**) K^+^-d[TG_4_T]_4_,*^31^* (**2**) K^+^-d[G_4_T_4_G_4_]2,*^32^* and (**3**) K+-d(G_4_T_4_)_3_G_4_.*^33^* Here, **1** is an all-*anti*-, parallel-stranded tetramer while **2** and **3** form antiparallel-stranded homodimers and monomers, respectively, with mixed *syn*-*/anti*-base conformations (Fig. 1B). The orientational differences between bases in stacked G-quartets of the model complexes are shown in detail in Fig. S1. All three complexes formed as expected based on their migration in native PAGE experiments (Fig. S2), with no indications of incomplete assembly or the formation of higher-order oligomerization states.

**Fig. 1.**
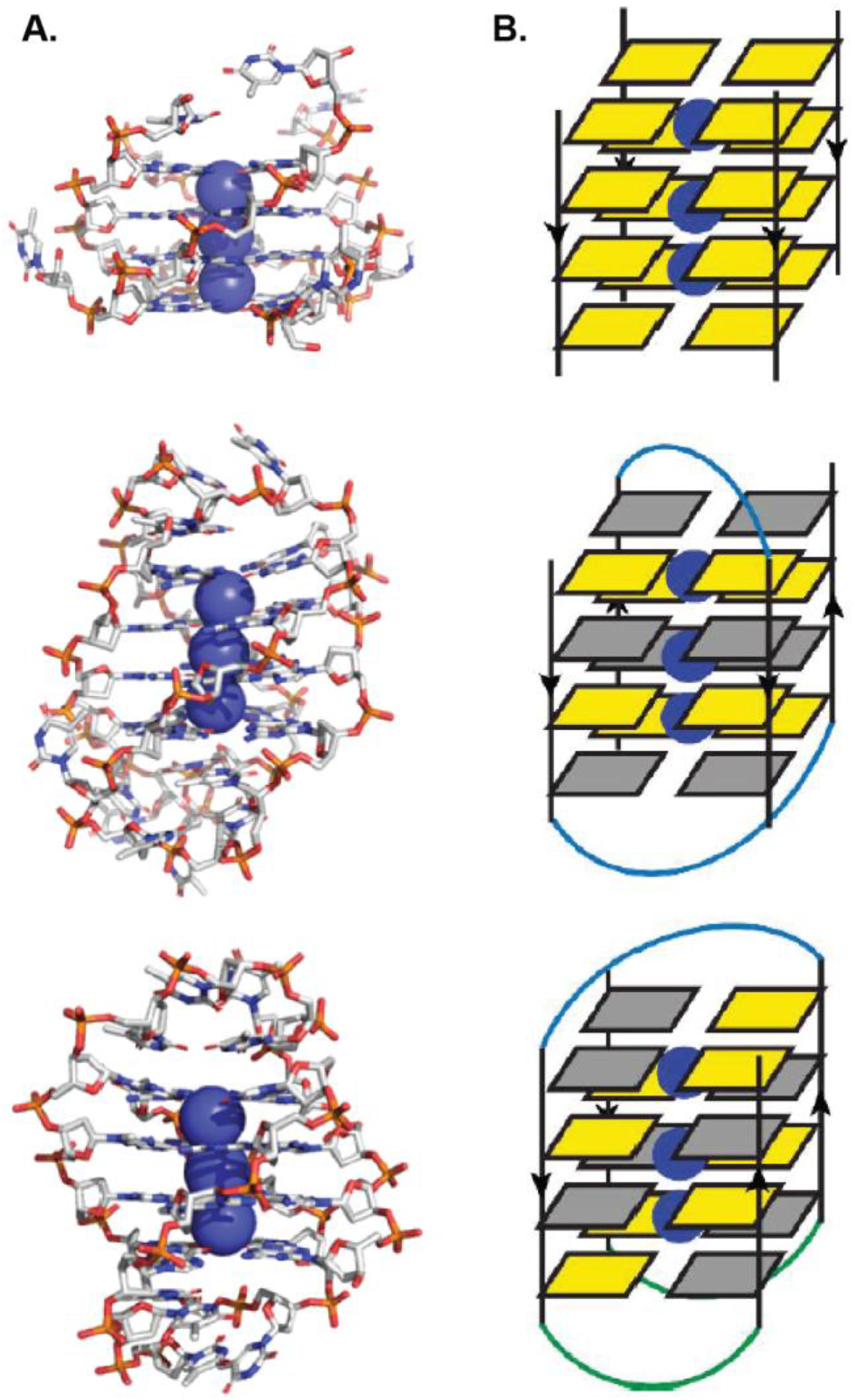
Parallel- and antiparallel-stranded G-quadruplexes. A. Structural models of K^+^-d[TG_4_T]_4_ (*top*, from PDB ID: 244D), K^+^-d[G_4_T_4_G_4_]_2_ (*middle*, from PDB ID: 2GWQ), and K^+^-d(G_4_T_4_)_3_G_4_ (*bottom*, from PDB ID: 201D). B. Schematic representation of the three complexes in (A) showing 5’-3’ strand orientations (arrows), *anti*-guanine bases (yellow) and *syn*-guanine bases (grey). Chemical structures and guanine conformations within the two central G-quartets of each complex are shown in Fig. S1.

In 2D IR spectra of **1** – **3** collected with parallel (*XXXX*) pump-probe polarization, the v(0-1) and v(1-2) features of guanine I, guanine II and thymine III modes are apparent along the diagonal and weak cross-peaks are observed (Fig. S3). The guanine III and guanine IV features below 1600 cm^−1^ are weak but are detected via cross-peaks (*vide infra*). The guanine C6=O (guanine I) frequencies are blue shifted compared to free guanosine monophosphate (GMP),*^27^* and the shift is larger in **1** than in **2** and **3**, due to quasi-parallel base orientation in the all-*anti*-structure, which enhances the inter-layer coupling effect.*^30^* To analyze the cross-peaks, we focus on those appearing below the diagonal (ω_probe_ > ω_pump_) because they are relatively free of interference.

### 3.2. Polarization dependent cross-peak enhancement in antiparallel-stranded G-quadruplexes

To examine polarization dependence, we collected 2D IR spectra in both *XXXX* (parallel) and *YYXX* (perpendicular) polarizations.*^34^* The spectra of **1** – **3** are compared in Fig. 2. For **1**, the intensities of the cross-peaks show only minor variations between *XXXX* (Fig. 2, left) and *YYXX* (Fig. 2, middle) spectra, which is clear from an overlay of the normalized vertical slices (Fig. 2, right). The absence of cross-peak enhancement in **1** mirrors the behavior of GMP (Fig. S4), suggesting that the symmetry of delocalized normal modes in the all-*anti*-structure does not differ substantially from that of the free guanine base. In contrast, both of the antiparallel-stranded G-quadruplexes (**2**, **3**) show a substantial increase in cross-peak intensity in *YYXX* spectra, particularly between guanine I (ω_pump_ ≈ 1670 cm^−1^) and guanine II (ω_pump_ ≈ 1600 cm^−1^) modes. We note that the polarization dependence of GMP cross-peaks is somewhat smaller than that observed by Peng *et al*.,*^29^* possibly due to solution conditions, differences between experimental setups, or both. Nevertheless, the method clearly distinguishes between parallel-stranded and antiparallel-stranded G-quadruplexes.

**Fig. 2.**
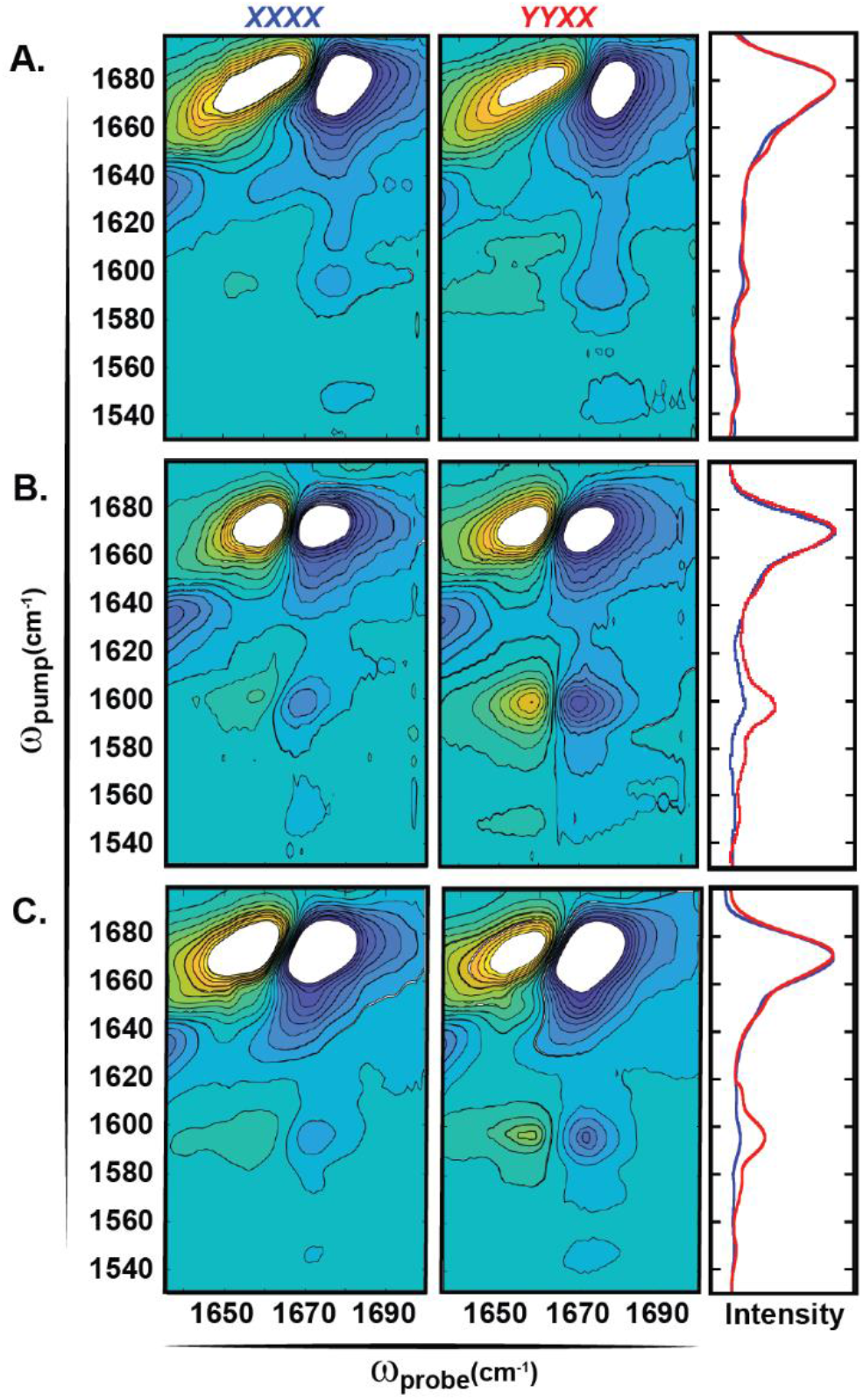
2D IR spectra of model G-quadruplexes. 2D IR spectra of (A) K^+^-d[TG_4_T]_4_, (B) K^+^-d[G_4_T_4_G_4_]_2_, and (C) K^+^-d(G_4_T_4_)_3_G_4_ collected in *XXXX* (left panels) and *YYXX* (center panels) polarizations. Spectra are scaled to 50% of the guanine I diagonal intensity. Overlays of vertical slices through the guanine I maxima and normalized to their respective maxima are shown in the right panels for *XXXX* (blue) and *YYXX* (red) spectra.

To further highlight the polarization dependence, we subtracted vertical slices of the *XXXX* spectra from the corresponding slice of the *YYXX* spectra of each model G-quadruplex (Fig. 3). For **1**, the *YYXX* - *XXXX* slice difference spectrum is within the level of noise. For **2** and **3**, the slice differences approach 20 – 30% of the guanine I diagonal intensity for the guanine I-II cross-peak (ω_pump_ ≈ 1600 cm^−1^) and 10 – 15% for the guanine I-III cross-peak (ω_pump_ ≈ 1580 cm^−1^). Interestingly, in **2** and **3** the latter differs between structures, suggesting that distinguishing the topologies of different antiparallel-stranded G-quadruplexes may become possible with the aid of quantum mechanical calculations and polarization angle scanning.*^28,30,34,35^*

**Fig. 3.**
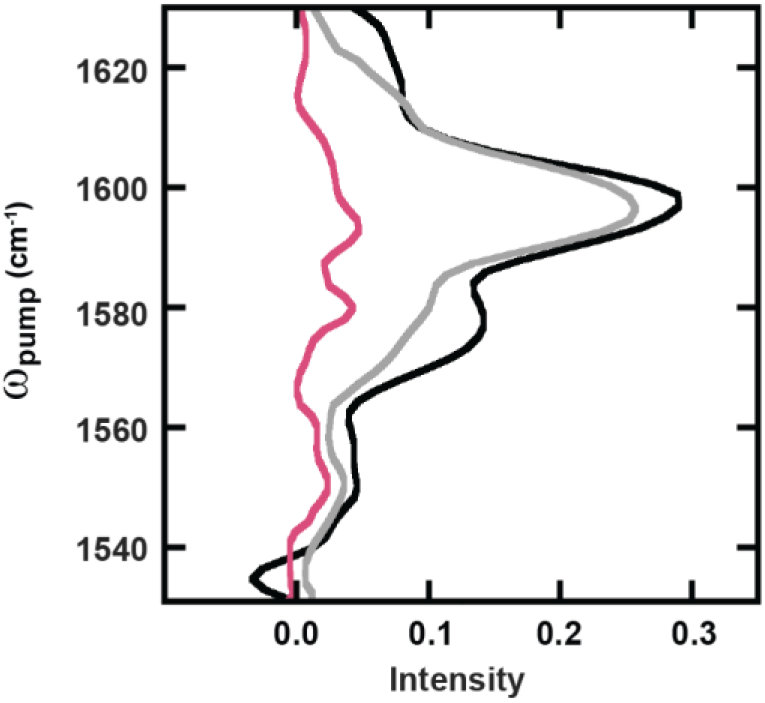
Polarization-dependence of cross-peaks. Slice difference spectra (*YYXX* - *XXXX*) in the cross-peak region at the guanine I maxima along ω_probe_ for K^+^-d[TG_4_T]_4_ (red), K^+^-d[G_4_T_4_G_4_]_2_ (black), and K^+^-d(G_4_T_4_)_3_G_4_ (grey).

### 3.3. Application to the structural analysis of a Bcl-2 promoter variant

Having established this spectral marker of strand orientation in G-quadruplexes, we tested our method on a 30-nucleotide sequence derived from the Bcl-2 promoter. In previous work, the variant Pu30_3T4AA was found to adopt an all-*anti*-, parallel-stranded G-quadruplex with a 1:13:1 chain reversal loop configuration using a combination of NMR spectroscopy, UV-CD spectroscopy, and chemical footprinting (Fig. 4A).*^36^* In the FTIR spectrum of Pu30_3T4AA prepared with 140 mM K^+^, four peaks are resolved between 1500 – 1700 cm^−1^ (Fig. 4B). These arise from mixtures of signals from the constituent bases, and it is known from previous work that the ~1670 cm^−1^ feature contains contributions from guanine I modes, the ~1625 cm^−1^ feature contains contributions from thymine modes, and the ~1590 cm^−1^ feature contains contributions from guanine II modes.*^27,29^*

**Fig. 4.**
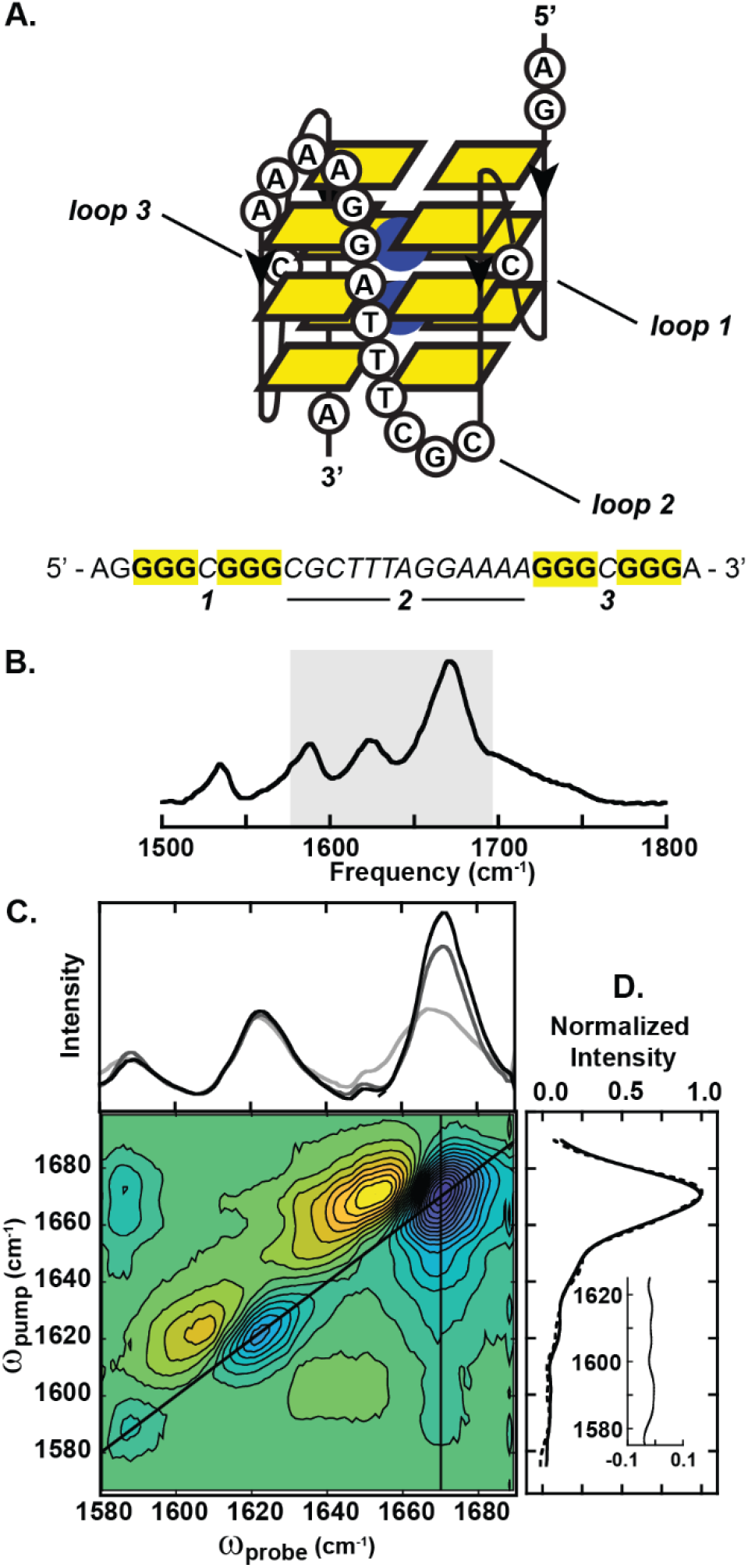
Analysis of the Bcl-2 promoter variant. A. *Top:* Model structure of Pu30_3T4AA showing locations of three loops (adapted from Ref. 36). *Bottom:* Sequence of Pu30_3T4AA showing proposed G-quadruplex forming guanine tracts (bold, yellow) and locations of loops 1 – 3 (italics). B. FTIR spectrum of Pu30_3T4AA in the presence of 140 mM K^+^. The region shaded in grey is detected in 2D IR spectra. C. 2D IR spectroscopy of Pu30_3T4AA. *Top:* diagonal slices of spectra collected in the presence of 0 mM (light grey), 40 mM (dark grey), and 140 mM (black) K^+^ ions. *Bottom: YYXX* spectrum of Pu30_3T4AA showing positions of diagonal and vertical slices. D. Overlay of *XXXX* (solid) and *YYXX* (dashed) vertical slices (140 mM K^+^). *Inset: YYXX* - *XXXX* slice difference. Additional spectra used in this analysis are shown in Fig. S5.

Diagonal slices through the *XXXX* 2D IR spectra of Pu30_3T4AA prepared with 0 mM, 40 mM, and 140 mM K^+^ (Fig. 4C, *top*) indicate narrowing of the high-frequency (guanine I) feature and a concomitant blue shift consistent with the assembly of a G-quadruplex.*^12,27^* Notably, the shift is complete at 40 mM K^+^, suggesting that the saturation threshold for monomeric G-quadruplexes is lower than that for tetramers.*^27^* Minor perturbation in the guanine II region is also observed, but the ~1625 cm^−1^ feature is not K^+^-dependent, indicating that adenine, cytosine, and thymine bases absorbing in the region are excluded from the K^+^-assembled array. In the *YYXX* 2D IR spectrum (Fig. 4C, *bottom*) guanine I-II cross-peaks appear above and below the diagonal. Vertical slices through the guanine I maximum (ω_probe_ = 1671 cm^−1^) in *YYXX* and *XXXX* spectra (Fig. 4D) show no polarization dependent enhancement of cross-peaks, similar to model complex **1** (Fig. 2A).

The 2D IR spectra of the Bcl-2 promoter variant Pu30_3T4AA provide a wealth of structural information. First, only guanine modes are K^+^-dependent, so all other regions of the sequence remain disordered (e.g., within loops) upon G-quadruplex assembly. Second, the slice difference spectra indicate all-*anti*-guanine base conformations consistent with the parallel-stranded structure. Third, after confirming the parallel strand orientations, it is possible to use the frequency of the bright high-frequency guanine I mode at K^+^ saturation as a measure of the motif size; the observed guanine I frequency of 1671 cm^−1^ is nearly identical to that of K^+^-d[TG_3_T]_4_, indicating that the ordered G-quadruplex is composed of three stacked G-quartets.*^27^* Together, these observations capture the major features of the NMR-based structure*^36^* and only the global topology, which is dependent on the organization of loops, is unknown from IR analysis. In the absence of the reference structure, the parameters defined by 2D IR narrow the plausible set of folds, and molecular dynamics simulations could be employed to examine their stabilities.*^37^* Site-specific isotope labeling could also be used to confirm which guanines exists within the G-quadruplex core via metal-dependent frequency shifts and exciton disruption.*^27^*

## 4. Conclusions

These results demonstrate that the analysis of vibrational coupling among G-quartets via the polarization dependence of 2D IR cross-peaks is sufficient to distinguish between parallel-stranded and antiparallel-stranded structures. Our method, which only requires the comparison to two 2D IR spectra, reveals this critical structural feature rapidly and with minimal sample requirements. Where parallel-stranded structures are identified, previously defined guanine I frequency trends can be used to define the size of the ordered motif. The insights gained from 2D IR alone are already competitive with other low- to medium-resolution techniques for G-quadruplex characterization.*^7–10^* However, because stacking-dependent frequency trends in antiparallel-stranded systems remain uncharacterized, the full analysis is currently limited to the subset of parallel-stranded systems. Future experiments using site-specific isotope labeling may be applied to probe pairwise coupling between bases and exciton disruption among mixed *syn*-/*anti*-G-quartet layers, broadening the scope of the approach. With the aid of quantum mechanical calculations and the independent polarization control of pulses, additional features of the spectra, such as cross-peaks in the guanine II-IV region, may be assigned to provide insight into challenging structures.*^28–30,35,38^* Importantly, guanine I modes are well-resolved in 2D IR spectra on account of the fourth-power dependence of signals on transition dipole moments,*^13^* allowing our method to be applied to G-quadruplexes with complex loops and flanking sequences. Moreover, 2D IR is largely insensitive to sample morphology, and can be used to analyze G-quadruplexes formed within disordered or insoluble structures formed by longer sequences. Thus, the analysis described here is not only useful for structure determination but can also be applied to screen G-quadruplex variants across a broad range of conditions to optimize the design of selective molecular probes or drug candidates.*^39,40^* With continued improvements in experimental and theoretical methods, we expect 2D IR spectroscopy to become a key tool in efforts to characterize and understand these important biological structures.

## Supplementary Material

Supplementary Information to this article includes a table of oligonucleotide sequences, structural models of G-quadruplexes, native polyacrylamide gel electrophoresis (PAGE) results, and 2D IR spectra as described in the text.

## References

[1] Varshney, D., Spiegel, J., Zyner, K., Tannahill, D., and Balasubramanian, S. (2020) The regulation and functions of DNA and RNA G-quadruplexes, Nat. Rev. Mol. Cell Biol. 21, 459–474.

[2] Simone, R., Fratta, P., Neidle, S., Parkinson, G. N., and Isaacs, A. M. (2015) G-quadruplexes: Emerging roles in neurodegenerative diseases and the non-coding transcriptome, FEBS Lett. 589, 1653–1668.

[3] Bochman, M. L., Paeschke, K., and Zakian, V. A. (2012) DNA secondary structures: stability and function of G-quadruplex structures, Nat. Rev. Genet. 13, 770–780.

[4] Burge, S., Parkinson, G. N., Hazel, P., Todd, A. K., and Neidle, S. (2006) Quadruplex DNA: sequence, topology and structure, Nucleic Acids Res. 34, 5402–5415.

[5] Asamitsu, S., Obata, S., Yu, Z., Bando, T., and Sugiyama, H. (2019) Recent Progress of Targeted G-Quadruplex-Preferred Ligands Toward Cancer Therapy, Molecules 24.

[6] Ma, D.-L., Zhang, Z., Wang, M., Lu, L., Zhong, H.-J., and Leung, C.-H. (2015) Recent Developments in G-Quadruplex Probes, Chem. Biol. 22, 812–828.

[7] Murat, P., Singh, Y., and Defrancq, E. (2011) Methods for investigating G-quadruplex DNA/ligand interactions, Chem. Soc. Rev. 40, 5293–5307.

[8] Reddy Sannapureddi, R. K., Mohanty, M. K., Gautam, A. K., and Sathyamoorthy, B. (2020) Characterization of DNA G-quadruplex Topologies with NMR Chemical Shifts, J. Phys. Chem. Lett. 11, 10016–10022.

[9] del Villar-Guerra, R., Trent, J. O., and Chaires, J. B. (2018) G-Quadruplex Secondary Structure Obtained from Circular Dichroism Spectroscopy, Angew. Chem. Int. Ed. 57, 7171–7175.

[10] Hu, M.-H., and Chen, X. (2020) New fluorescent light-up quinoxalines differentiate between parallel and nonparallel G-quadruplex topologies using different excitation/emission channels, Chem. Commun. 56, 4168–4171.

[11] Marchand, A., and Gabelica, V. (2016) Folding and misfolding pathways of G-quadruplex DNA, Nucleic Acids Res. 44, 10999–11012.

[12] Guzmán, M. R., Liquier, J., Brahmachari, S. K., and Taillandier, E. (2006) Characterization of parallel and antiparallel G-tetraplex structures by vibrational spectroscopy, Spectrochim. Acta A Mol. Biomol. Spectrosc. 64, 495–503.

[13] Hamm, P., and Zanni, M. (2011) Concepts and Methods of 2D Infrared Spectroscopy, Cambridge University Press, Cambridge.

[14] Le Sueur, A. L., Horness, R. E., and Thielges, M. C. (2015) Applications of two-dimensional infrared spectroscopy, Analyst 140, 4336–4349.

[15] Ganim, Z., Chung, H. S., Smith, A. W., DeFlores, L. P., Jones, K. C., and Tokmakoff, A. (2008) Amide I Two-Dimensional Infrared Spectroscopy of Proteins, Acc. Chem. Res. 41, 432–441.

[16] Ghosh, A., Ostrander, J. S., and Zanni, M. T. (2017) Watching Proteins Wiggle: Mapping Structures with Two-Dimensional Infrared Spectroscopy, Chem. Rev. 177, 10726–10759.

[17] Kim, Y. S., and Hochstrasser, R. M. (2009) Applications of 2D IR Spectroscopy to Peptides, Proteins, and Hydrogen-Bond Dynamics, J. Phys. Chem. B 113, 8231–8251.

[18] Cheatum, C. M., Tokmakoff, A., and Knoester, J. (2004) Signatures of β-sheet secondary structures in linear and two-dimensional infrared spectroscopy, J. Chem. Phys. 120, 8201–8215.

[19] Lomont, J. P., Ostrander, J. S., Ho, J.-J., Petti, M. K., and Zanni, M. T. (2017) Not All β-Sheets Are the Same: Amyloid Infrared Spectra, Transition Dipole Strengths, and Couplings Investigated by 2D IR Spectroscopy, J. Phys. Chem. B 121, 8935–8945.

[20] Lomont, J. P., Rich, K. L., Maj, M., Ho, J.-J., Ostrander, J. S., and Zanni, M. T. (2018) Spectroscopic Signature for Stable β-Amyloid Fibrils versus β-Sheet-Rich Oligomers, J. Phys. Chem. B 122, 144–153.

[21] Londergan, C. H., Wang, J., Axelsen, P. H., and Hochstrasser, R. M. (2006) Two-Dimensional Infrared Spectroscopy Displays Signatures of Structural Ordering in Peptide Aggregates, Biophys. J. 90, 4672–4685.

[22] Strasfeld, D. B., Ling, Y. L., Gupta, R., Raleigh, D. P., and Zanni, M. T. (2009) Strategies for Extracting Structural Information from 2D IR Spectroscopy of Amyloid: Application to Islet Amyloid Polypeptide, J. Phys. Chem. B 113, 15679–15691.

[23] Bruening, E. M., Schauss, J., Siebert, T., Fingerhut, B. P., and Elsaesser, T. (2018) Vibrational Dynamics and Couplings of the Hydrated RNA Backbone: A Two-Dimensional Infrared Study, J. Phys. Chem. Lett. 9, 583–587.

[24] Schauss, J., Kundu, A., Fingerhut, B. P., and Elsaesser, T. (2019) Contact Ion Pairs of Phosphate Groups in Water: Two-Dimensional Infrared Spectroscopy of Dimethyl Phosphate and ab Initio Simulations, J. Phys. Chem. Lett. 10, 6281–6286.

[25] Yang, M., Szyc, Ł., and Elsaesser, T. (2011) Femtosecond Two-Dimensional Infrared Spectroscopy of Adenine-Thymine Base Pairs in DNA Oligomers, J. Phys. Chem. B 115, 1262–1267.

[26] Hithell, G., Ramakers, L. A. I., Burley, G. A., and Hunt, N. T. (2018) Chapter 3 - Applications of 2D-IR Spectroscopy to Probe the Structural Dynamics of DNA, In Frontiers and Advances in Molecular Spectroscopy (Laane, J., Ed.), pp 77–100, Elsevier.

[27] Price, D. A., Kartje, Z. J., Hughes, J. A., Hill, T. D., Loth, T. M., Watts, J. K., Gagnon, K. T., and Moran, S. D. (2020) Infrared Spectroscopy Reveals the Preferred Motif Size and Local Disorder in Parallel Stranded DNA G-Quadruplexes, ChemBioChem 21, 2792–2804.

[28] Krummel, A. T., and Zanni, M. T. (2006) DNA Vibrational Coupling Revealed with Two Dimensional Infrared Spectroscopy: Insight into Why Vibrational Spectroscopy Is Sensitive to DNA Structure, J. Phys. Chem. B 110, 13991–14000.

[29] Peng, C. S., Jones, K. C., and Tokmakoff, A. (2011) Anharmonic Vibrational Modes of Nucleic Acid Bases Revealed by 2D IR Spectroscopy, J. Am. Chem. Soc. 133, 15650–15660.

[30] Jiang, Y., and Wang, L. (2020) Modeling the vibrational couplings of nucleobases, J. Chem. Phys. 152, 084114.

[31] Laughlan, G., Murchie, A. I., Norman, D. G., Moore, M. H., Moody, P. C., Lilley, D. M., and Luisi, B. (1994) The high-resolution crystal structure of a parallel-stranded guanine tetraplex, Science 265, 520.

[32] Schultze, P., Smith, F. W., and Feigon, J. (1994) Refined solution structure of the dimeric quadruplex formed from the *Oxytricha* telomeric oligonucleotide d(GGGGTTTTGGGG), Structure 2, 221–233.

[33] Wang, Y., and Patel, D. J. (1995) Solution Structure of the *Oxytricha* Telomeric Repeat d[G4(T4G4)3] G-tetraplex, J. Mol. Biol. 251, 76–94.

[34] Woutersen, S., and Hamm, P. (2000) Structure Determination of Trialanine in Water Using Polarization Sensitive Two-Dimensional Vibrational Spectroscopy, J. Phys. Chem. B 104, 11316–11320.

[35] Choi, J.-H., and Cho, M. (2010) Polarization-angle-scanning two-dimensional infrared spectroscopy of antiparallel β-sheet polypeptide: Additional dimensions in two-dimensional optical spectroscopy, J. Chem. Phys. 133, 241102.

[36] Agrawal, P., Lin, C., Mathad, R. I., Carver, M., and Yang, D. (2014) The Major G-Quadruplex Formed in the Human BCL-2 Proximal Promoter Adopts a Parallel Structure with a 13-nt Loop in K^+^ Solution, J. Am. Chem. Soc. 136, 1750–1753.

[37] Ortiz de Luzuriaga, I., Lopez, X., and Gil, A. (2021) Learning to Model G-Quadruplexes: Current Methods and Perspectives, Annu. Rev. Biophys. 50, 209–243.

[38] Lee, C., Park, K.-H., and Cho, M. (2006) Vibrational dynamics of DNA. I. Vibrational basis modes and couplings, J. Chem. Phys. 125, 114508.

[39] Ma, Y., Iida, K., and Nagasawa, K. (2020) Topologies of G-quadruplex: Biological functions and regulation by ligands, Biochem. Biophys. Res. Commun. 531, 3–17.

[40] Fritzsch, R., Donaldson, P. M., Greetham, G. M., Towrie, M., Parker, A. W., Baker, M. J., and Hunt, N. T. (2018) Rapid Screening of DNA–Ligand Complexes via 2D-IR Spectroscopy and ANOVA–PCA, Anal. Chem. 90, 2732–2740.

