## Supplementary Material for "The Polarization Dependence of 2D IR Cross-Peaks Distinguishes Parallel-Stranded and Antiparallel-Stranded DNA G-quadruplexes"

**Table S1. Sequences of model G-quadruplex forming oligonucleotides.**

| Complex | | Oligonucleotide Sequence |
|---|---|---|
| 1 | $K^+$-d[$TG_4T$]$_4$ | 5'-dT-dG-dG-dG-dG-dT-3' |
| 2 | $K^+$-d[$G_4T_4G_4$]$_2$ | 5'-dG-dG-dG-dG-dT-dT-dT-dT-dG-dG-dG-dG-3' |
| 3 | $K^+$-d($G_4T_4$)$_3G_4$ | 5'-dG-dG-dG-dG-dT-dT-dT-dT-dG-dG-dG-dG-dT-dT-dT-dT-dG-dG-dG-dG-dT-dT-dT-dT-dG-dG-dG-dG-3' |

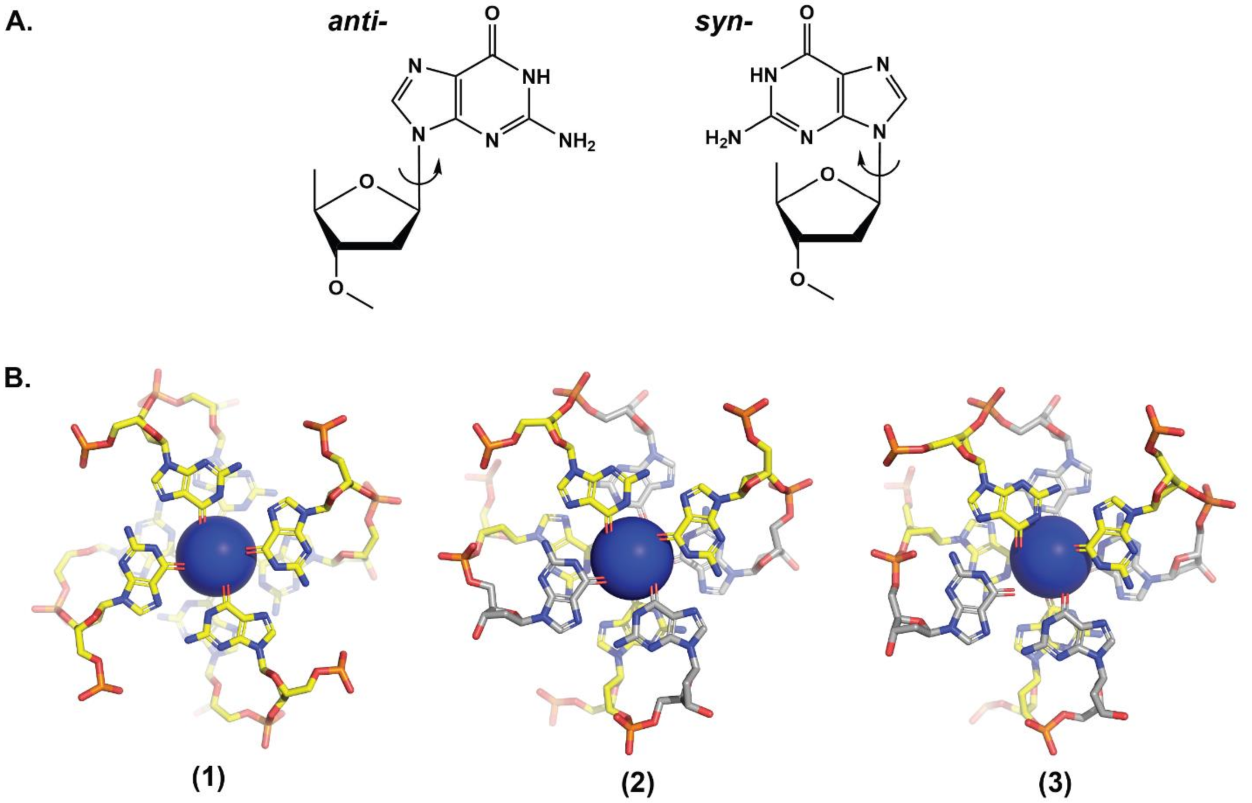


**Fig. S1. Base conformations in parallel- and antiparallel-stranded G-quadruplexes.** A. *anti*- (left) and *syn*- (right) conformations of the guanine base relative to the deoxyribose ring. Phosphodiester linkages are omitted for clarity. B. Two central G-quartets of (1) $K^+$-d[T$G_4$T]$_4$, (2) $K^+$-d[$G_4T_4G_4$]$_2$, and (3) $K^+$-d($G_4T_4$)$_3G_4$ are shown with $K^+$ (blue spheres), *anti*-guanines (yellow) and *syn*-guanines (grey). Structures were adapted from PDB IDs 244D (1), 2GWQ (2), and 201D (3) and rendered in PyMol (Schrodinger, New York, NY)

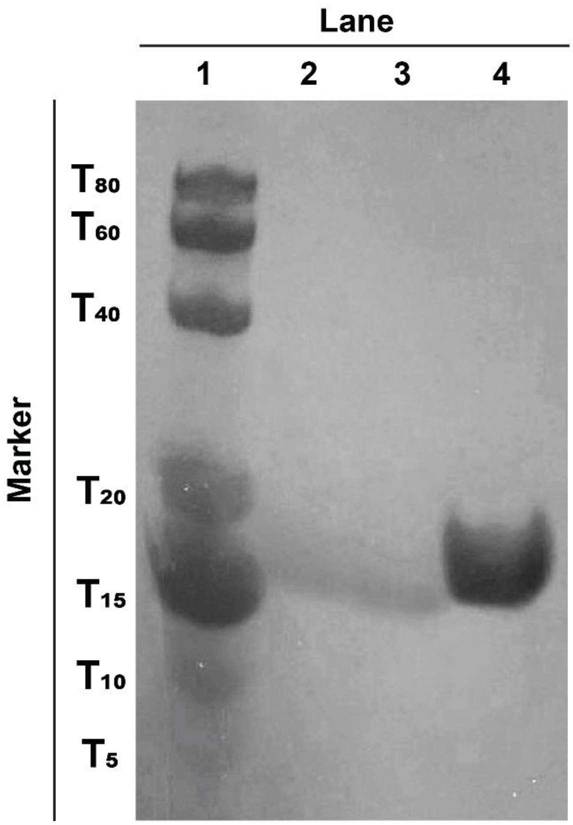


**Fig. S2. Native PAGE analysis of $K^+$-bound model G-quadruplexes.** Lane 1: Molecular weight marker composed of a mixture of polyT oligonucleotides. Lane 2: $[TG_4T]_4$. Lane 3: $[G_4T_4G_4]_2$. Lane 4: $(G_4T_4)_3G_4$.

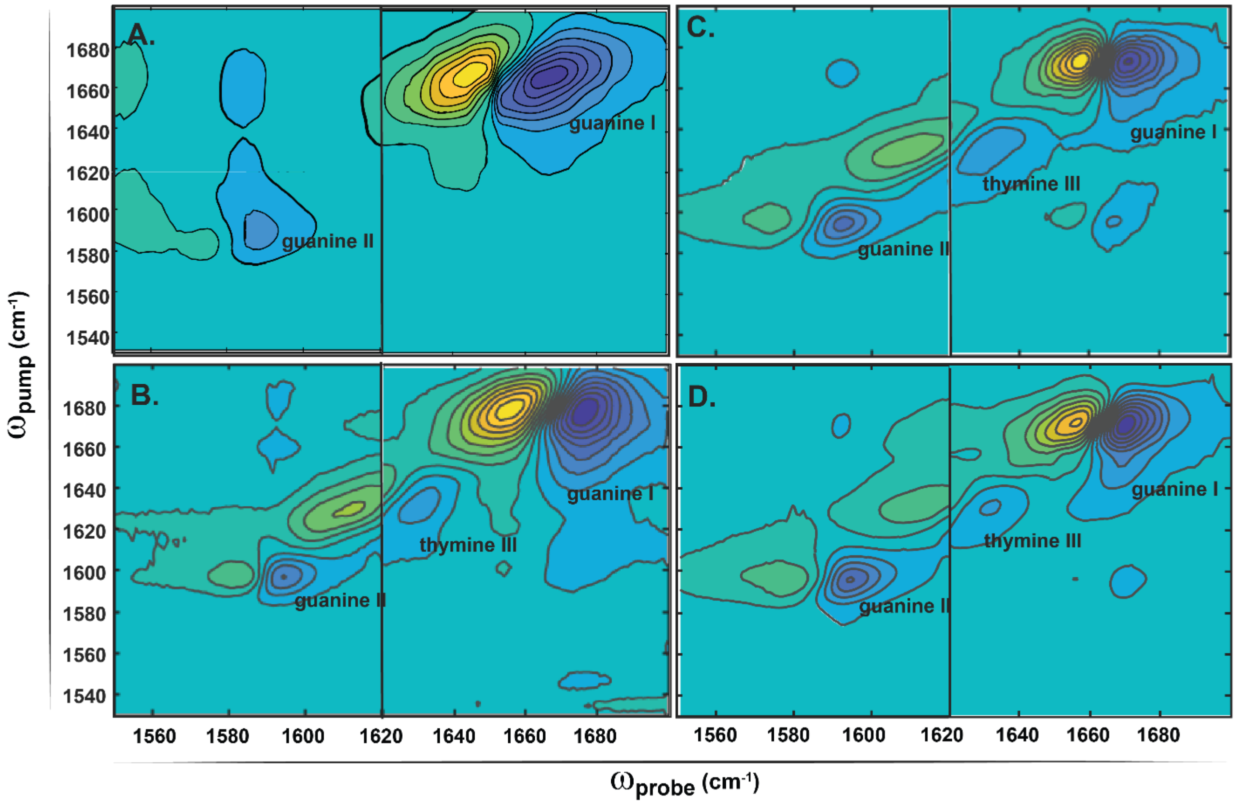


**Fig. S3. 2D IR spectra of GMP and model G-quadruplexes in *XXXX* polarization.** A. GMP. B. $K^+$-d[$TG_4T$]$_4$. C. $K^+$-d[$G_4T_4G_4$]$_2$. D. $K^+$-d($G_4T_4$)$_3G_4$. The GMP sample omitted $K^+$, and all G-quadruplex samples were prepared with 140 mM $K^+$. All spectra were collected at two overlapping monochromator positions to span the full range of guanine features and their v(1-2) transitions. Assigned peaks are labeled in the panels.

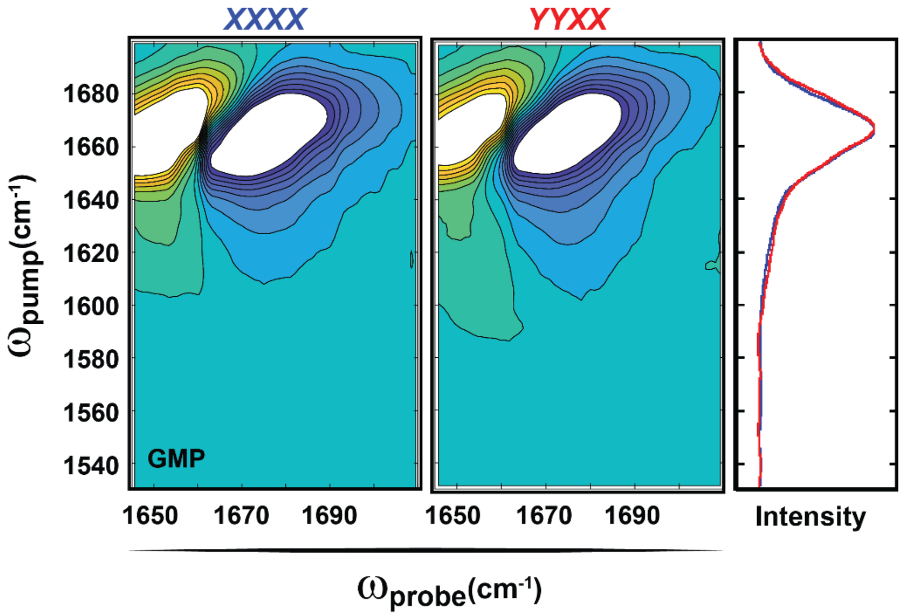


**Fig. S4. Polarization dependence of GMP cross-peaks.** 2D IR spectra of GMP in *XXXX* (left) and *YYXX* (middle) polarizations. Spectra are scaled to 50% of the guanine I diagonal ν(0-1) intensity. An overlay of normalized vertical slices through $\omega_{probe}$ = 1666 $cm^{-1}$ is shown in the right panel.

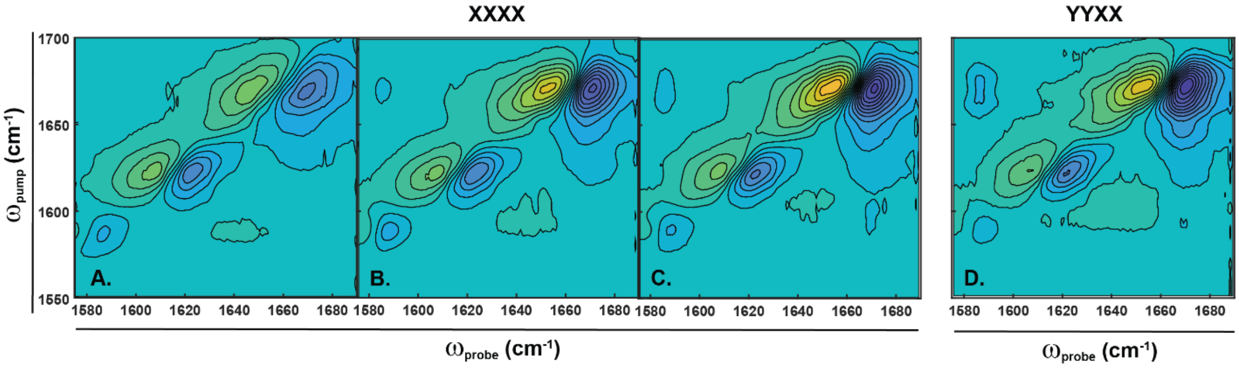


**Fig. S5. 2D IR spectra of Pu30_3T4AA.** 2D IR spectra of Pu30_3T4AA in *XXXX* polarization with (A) 0 mM, (B) 40 mM, and (C) 140 mM $K^+$. Spectra A-C are plotted with the same intensity scale to illustrate the changes in peak height. D. 2D IR spectrum of Pu30_3T4AA in *YYXX* polarization with 140 mM $K^+$.
